# The Influence of Knee Varus Deformity on the Kinematic and Dynamic Characteristics of Musculoskeletal Models During Gait

**DOI:** 10.1101/2023.11.02.565254

**Authors:** Sina Tabeiy, Morad Karimpour, Azizollah Shirvani, Arash Sherafat Vaziri

## Abstract

**Background:** Musculoskeletal modeling has paved the way of measuring kinematic and kinetic variables during motions. Nonetheless, since the commonly-used generic models are created based on averaged data; thus, they cannot accurately mimic subjects with skeletal deformities. To overcome this obstacle, one can build personalized models based on subject’s MRI or CT scan data, which is both time and money consuming. The other promising way is to manipulate generic models and create semi-personalized models to match with the individual’s skeletal system at the joint of interest.

**Research Question:** Can a semi-personalized model reduce marker error in gait analysis? How a semi-personalized model differentiates the ROM of the lower limb joints and muscle activation pattern while having varus deformity?

**Method:** We developed the varus-valgus tool (freely available on: https://simtk.org/projects/var-val-tool) in MATLAB using OpenSim Application Programming Interface (API) to incorporate varus-valgus deformity in the generic OpenSim models. A 36-year-old female subject with a complaint of knee pain participated in our study. The subject had 6.5 and 11.9 degrees of varus in the right and left leg, respectively. A semi-personalized model of the subject was first created using generic OpenSim models. Then, markers’ error during Inverse Kinematic (IK), joints Range of Motion (ROM) and the activation of Tensor Fasciae Latae (TFL), a knee adductor, and Gracilis, a knee abductor, were calculated and compared between a semi-personalized model and a generic model.

**Results:** Significant difference was observed in markers’ error during IK between generic and semi-personalized models (p<0.05). Substantial alterations were found in the ROM of the hip, knee and ankle joints while using semi-personalized model. Moreover, the activation pattern of TFL experienced a dramatic rise whereas Gracilis saw a fall during each gait cycle in semi-personalized models.

**Significance:** Implementing varus-valgus deformity in the generic models substantially reduces markers’ error which leads to more accurate results. It was observed that semi-personalized models showed different ROM compared to generic ones.

## Introduction

Motion capture (MoCap) systems have enabled researchers to investigate joints’ function during simple to sophisticated movements (1, 2). Among all trials captured in MoCap laboratories, the gait trial is known as a standard criteria to estimate the kinetics and kinematics of the body joints. The outcomes of MoCap systems are invaluable since they can be used in clinical assessments, injury prevention, and immediate evaluation of treatment (3, 4).

One way to analyze the data collected from a gait laboratory is to utilize musculoskeletal models in simulator software, like OpenSim (5, 6) or AnyBody (7). OpenSim is more favorable thanks to the ability to perform inverse methods (8), which allows researchers to conduct additional analyses, including the calculation of Joint Reaction Force (9, 10), muscle-tendon length (11), muscle-tendon force (12), muscle moment arm (13). To do so, OpenSim uses generic musculoskeletal models which have been created based on the data from healthy or cadaveric musculoskeletal systems (2). Generic models are user-friendly and approximately accurate in estimating joint kinematics and kinetics compared to in vivo methods (2, 9, 14-17). Nevertheless, since they are created based on measurements from healthy subjects, using them to analyze subjects with skeletal deformities, such as genu varum and genu valgum, may result in an inaccurate outcome.

Genu varum and genu valgum, also known as knee varus and knee valgus, are the two common lower-extremity deformities in the frontal plane (18). Varus or valgus deformity could lead to uneven load distribution on the tibiofemoral joint (19-21), which may cause knee osteoarthritis (OA) in the long run (22, 23). The magnitude of the knee varus or knee valgus can be calculated through subjects’ CT scan, MRI, or X-ray images using the three orthopedic indices called mechanical Medial Proximal Tibial Angle (mMPTA), mechanical Lateral Distal Femoral Angle (mLDFA) and mechanical Femoral Tibial Angle (mFTA) (24, 25). Since in generic models, mMPTA and mLDFA are in their normal range, 90 degrees, analyzing patients with varus or valgus by generic models will cause imprecise results, and eventually, clinical misinterpretation.

To overcome this issue, patient-specific models have been used (2, 26), and until now, different methods of creating patient-specific models have been introduced (1, 8, 27-29). Although studies have proved that patient-specific models mimic the joint contact forces more accurately (2, 30), creating them is time-consuming and cost-intensive (27, 31). One possible way to overcome these problems is to manipulate generic models and incorporate varus or valgus deformity in models to match the subject’s musculoskeletal system.

To our best knowledge, no tool has been presented to easily and quickly modify varus or valgus angulation in OpenSim models. Therefore, this study aimed to (1) present a tool to manipulate varus or valgus angulations in generic OpenSim models, and for the first step (2) evaluate the presented tool using a subject with varus angulation.

## Methods

The varus-valgus tool (freely available on: https://simtk.org/projects/var-val-tool) is created by the application programming interface (API) of OpenSim 4.x in MATLAB (2021b; Math Works Inc., MA). This tool can modify OpenSim models to create semi-personalized models incorporating varus or valgus deformity (Figure 1). The varus-valgus tool creates a deformation pattern according to (1) the varus or valgus angulation based on the magnitude of mLDFA and mMPTA; and (2) the location of Center of Rotation Angulation (CORA) in the femur and tibia bone. The deformation pattern manipulates the musculoskeletal pathways and location of virtual markers to develop a semi-personalized (Figure 1).

**Figure 1).**
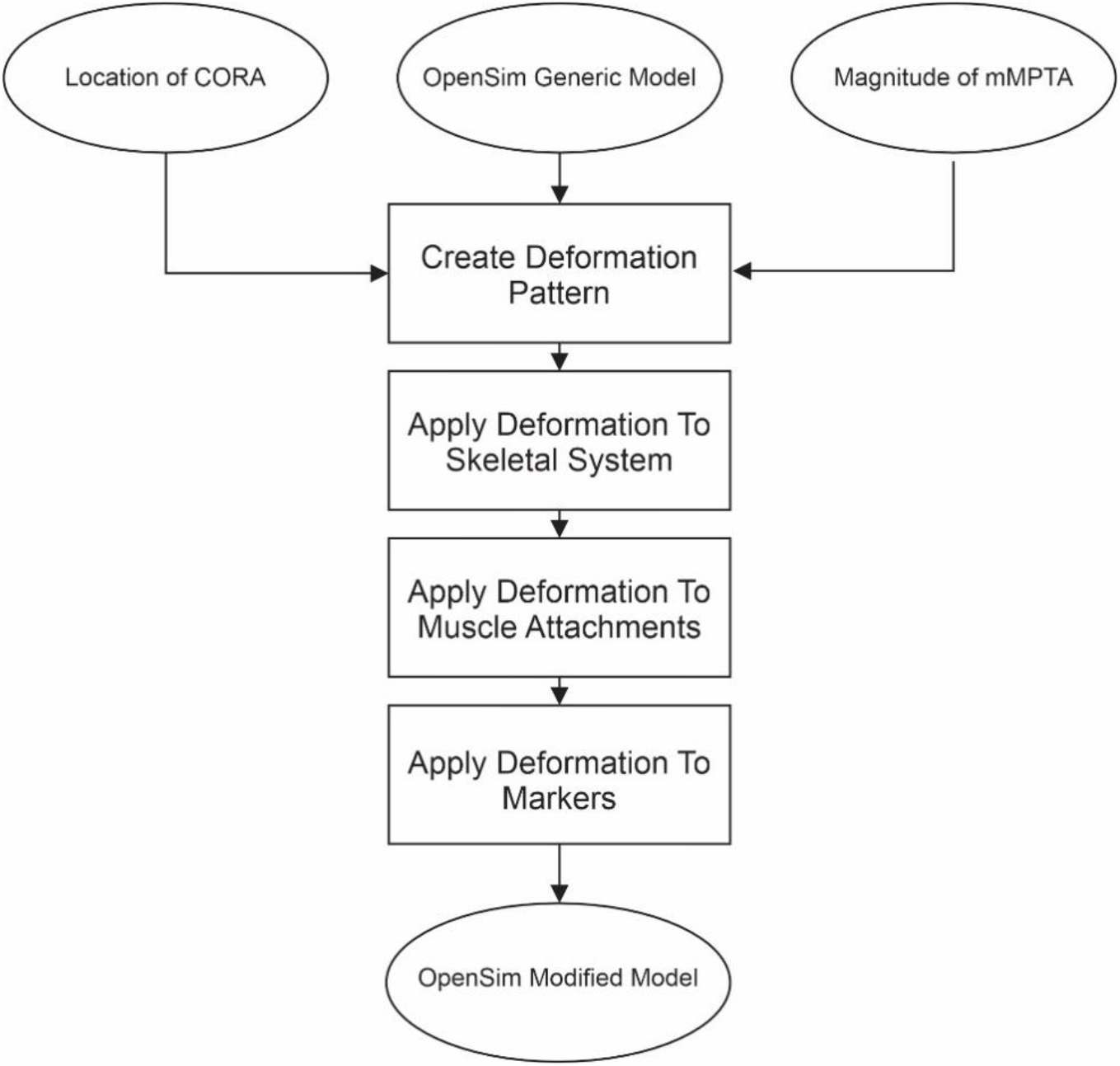
The flow chart of the steps needed to create a semi-personalized model using the varus-valgus tool.

**Figure 2).**
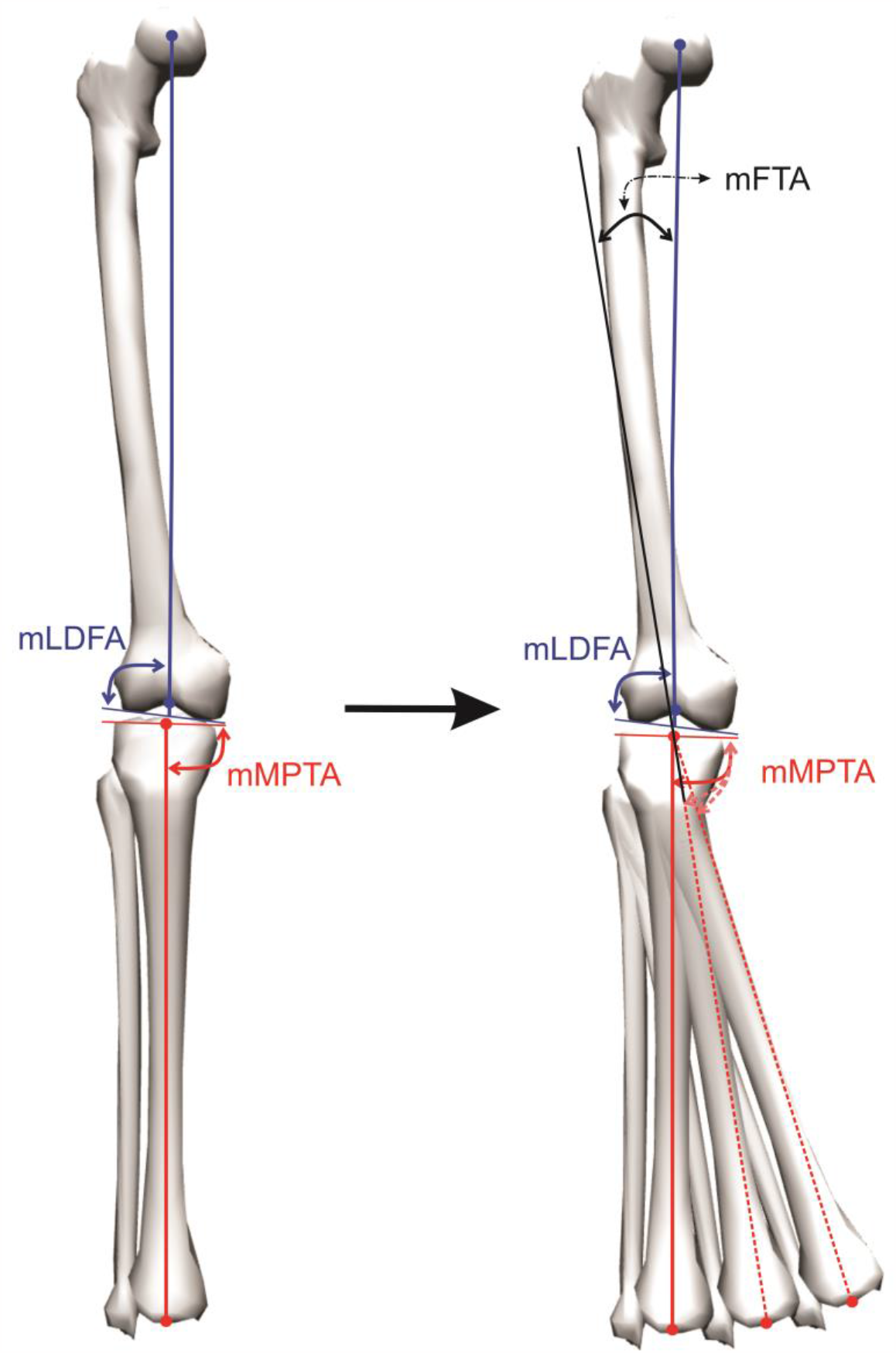
Schematic of m MPTA, mLDFA, and mFTA. The generic model is shown in the left side and semi-personalized models are in the right side.

A 36-year-old, otherwise healthy female subject with a complaint of knee pain participated in this study after giving informed consent before the experiments. This subject had mMPTA of 85.1 and mFTA of 6.5 degrees in varus in the right leg; mMPTA of 84.9 and mFTA of 11.9 degrees in varus in the left leg based on the assessment of her CT scan. Then, a semi-personalized model was developed using the varus-valgus tool to match the subject’s skeletal system.

After generating the semi-personalized model, the subject completed a normal gait trial at the gait laboratory of Djavad Mowafaghian Research Centre of Intelligent Neuro-Rehabilitation Technologies (Tehran, IR). Plug-in-gate marker set consisting of 39 reflective markers was used for this subject. Marker trajectories were recorded at 120 Hz using a ten-camera 3-dimensional motion capture system (MX13 cameras, Vicon Motion Systems, Oxford, UK). Ground Reaction Force (GRF) was captured using two force plates (AMTI, Watertown, MA, USA), synched with the motion capture system during the trial. The raw data collected from the gait laboratory was then labeled and filtered using Vicon Nexus 2.9.3 (Vicon Motion Systems, Oxford, UK).

To evaluate the varus-valgus tool, the models Rajagopal, Gait 2392, and Gait 2354 were semi-personalized in this study. The generic and semi-personalized models were scaled by the same scaling setup using equal weighting for markers and coordinates. Both models underwent Inverse Kinematics (IK) and Static Optimization (SO) with the exact weighting and cutoff frequency. Maximum and mean error of lower extremity markers during IK were plotted using a MATLAB code (https://github.com/johnjdavisiv/opensim-ik-errors). Then, normal Q-Q plots of the difference of each marker pair were then checked to follow a normal distribution. Finally, differences in the maximum error of lower extremity markers were compared by employing paired-samples T-tests (version 26, IBM SPSS Statistics, Chicago, IL, USA).

## Results

Maximum and mean error of each marker were calculated for both generic and semi-personalized models during IK (Figure 3). The markers’ error significantly reduced in semi-personalized Gait 2354 (Table 1 and Figure 3), and semi-personalized Rajagopal model (Table 2 and Figure 4).

**Table 1).** Result of paired-samples t-tests for each marker pair of generic and semi-personalized Gait 2354 model

|  | Paired Differences |  |  |  |  | t | df | Sig. (2-tailed) |
| --- | --- | --- | --- | --- | --- | --- | --- | --- |
|  | Mean | Std. Deviation | Std. Error Mean | 95% Confidence Interval of the Difference |  |  |  |  |
|  |  |  |  | Lower | Upper |  |  |  |
| LANK | 0.0056 | 0.0016 | 0.0001 | 0.0054 | 0.0059 | 41.1498 | 137.0000 | 0.0000 |
| LASI | 0.0050 | 0.0017 | 0.0001 | 0.0047 | 0.0053 | 34.1792 | 137.0000 | 0.0000 |
| LHEE | 0.0000 | 0.0052 | 0.0004 | -0.0009 | 0.0008 | -0.0672 | 137.0000 | 0.9465 |
| LKNE | 0.0068 | 0.0011 | 0.0001 | 0.0066 | 0.0070 | 73.3148 | 137.0000 | 0.0000 |
| LTOE | 0.0042 | 0.0020 | 0.0002 | 0.0039 | 0.0045 | 25.2077 | 137.0000 | 0.0000 |
| RANK | 0.0062 | 0.0013 | 0.0001 | 0.0060 | 0.0065 | 54.8373 | 137.0000 | 0.0000 |
| RASI | 0.0053 | 0.0016 | 0.0001 | 0.0050 | 0.0055 | 39.4629 | 137.0000 | 0.0000 |
| RHEE | 0.0046 | 0.0024 | 0.0002 | 0.0042 | 0.0050 | 22.5445 | 137.0000 | 0.0000 |
| RKNE | 0.0067 | 0.0005 | 0.0000 | 0.0066 | 0.0068 | 147.8157 | 137.0000 | 0.0000 |
| RTOE | 0.0058 | 0.0012 | 0.0001 | 0.0056 | 0.0060 | 55.8712 | 137.0000 | 0.0000 |

**Table 2).** Result of paired-samples t-tests for each marker pair of generic and semi-personalized Rajagopal model

|  | Paired Differences |  |  |  |  | t | df | Sig. (2-tailed) |
| --- | --- | --- | --- | --- | --- | --- | --- | --- |
|  | Mean | Std. Deviation | Std. Error Mean | 95% Confidence Interval of the Difference |  |  |  |  |
|  |  |  |  | Lower | Upper |  |  |  |
| LANK | 0.0035 | 0.0008 | 0.0001 | 0.0033 | 0.0036 | 48.9276 | 137.0000 | 0.0000 |
| LASI | 0.0032 | 0.0016 | 0.0001 | 0.0030 | 0.0035 | 23.3820 | 137.0000 | 0.0000 |
| LHEE | 0.0099 | 0.0038 | 0.0003 | 0.0092 | 0.0105 | 30.0900 | 137.0000 | 0.0000 |
| LKNE | 0.0038 | 0.0042 | 0.0004 | 0.0030 | 0.0045 | 10.4828 | 137.0000 | 0.0000 |
| LTOE | 0.0061 | 0.0046 | 0.0004 | 0.0053 | 0.0069 | 15.5119 | 137.0000 | 0.0000 |
| RANK | 0.0012 | 0.0008 | 0.0001 | 0.0011 | 0.0013 | 17.1040 | 137.0000 | 0.0000 |
| RASI | 0.0027 | 0.0017 | 0.0001 | 0.0024 | 0.0030 | 19.2282 | 137.0000 | 0.0000 |
| RHEE | 0.0007 | 0.0010 | 0.0001 | 0.0005 | 0.0009 | 7.8889 | 137.0000 | 0.0000 |
| RKNE | 0.0029 | 0.0013 | 0.0001 | 0.0027 | 0.0031 | 26.6089 | 137.0000 | 0.0000 |
| RTOE | 0.0033 | 0.0030 | 0.0003 | 0.0028 | 0.0038 | 12.8643 | 137.0000 | 0.0000 |

**Figure 3).**
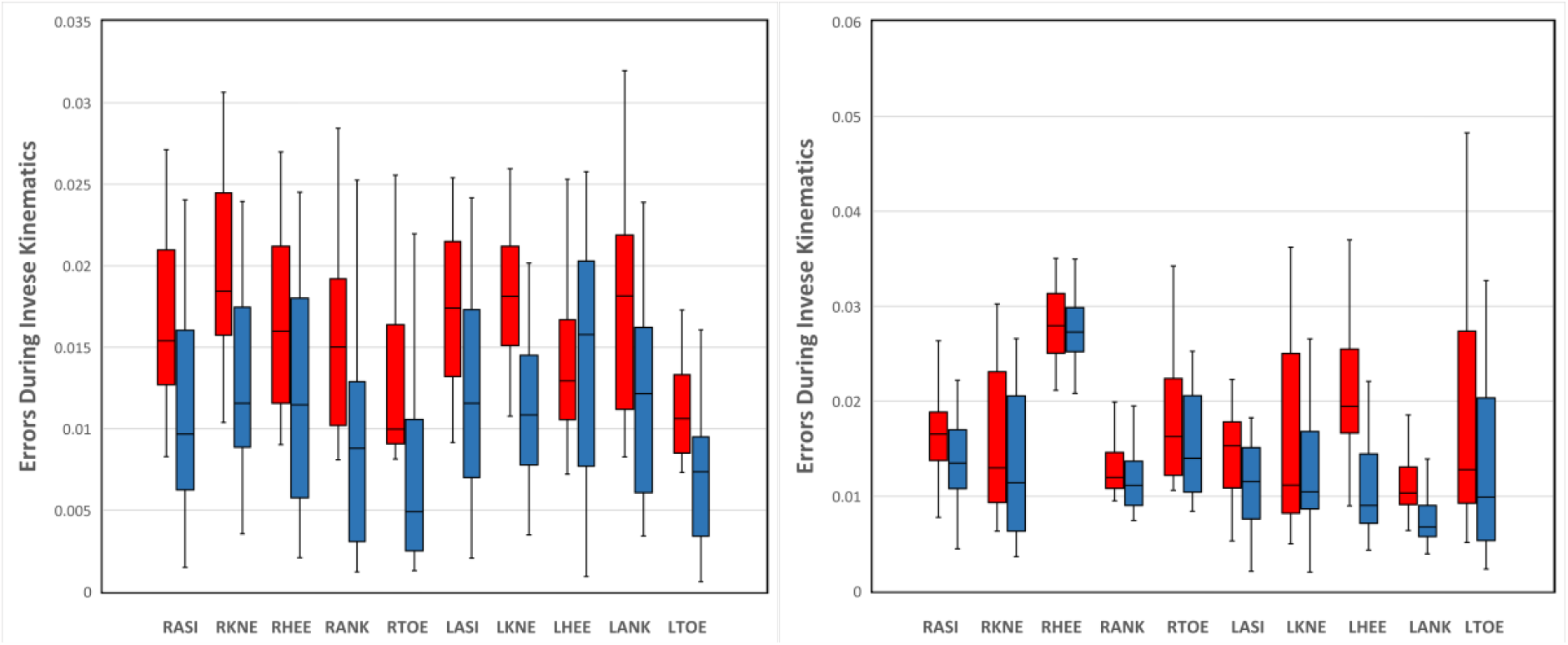
Errors during Inverse Kinematic for all markers using the Gait 2354 (left) and Rajagopal model (right); the blue boxes indicate markers’ errors of the semi-personalized model, and the red ones indicate the generic model.

**Figure 4).**
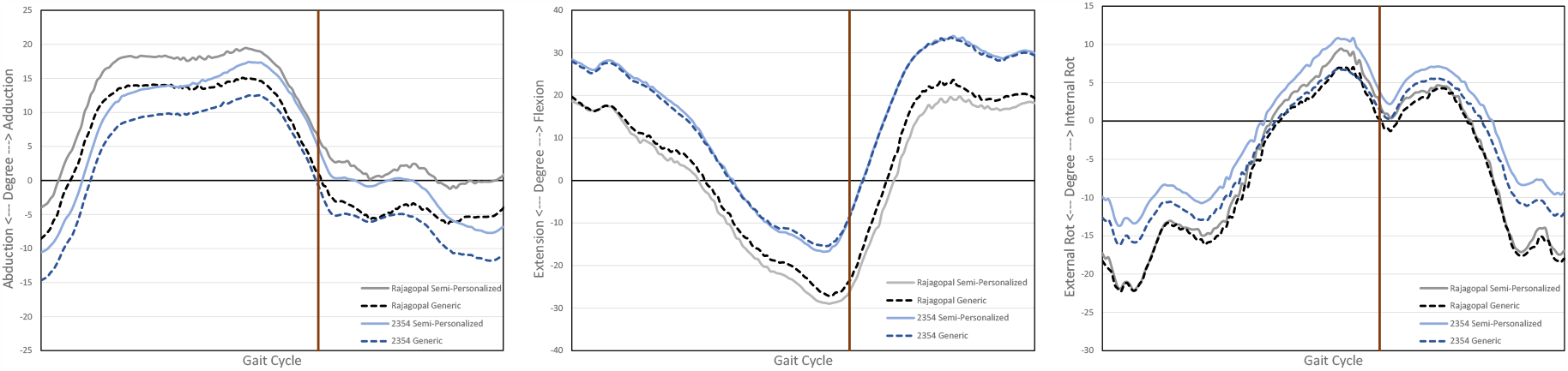
Hip abduction/adduction, flexion/extension, and internal/external rotation during a complete gait cycle.

IK results of hip joint movements are depicted in the figures below (Figure 4, 5, and 6). Although apparent changes can be seen in all three hip movements, the difference in hip adduction (Figure 4) is more evident than the other movements (Figures 5 and 6). IK results for knee flexion/extension and ankle dorsi/plantar flexion are also shown in Figure 7.

**Figure 5).**
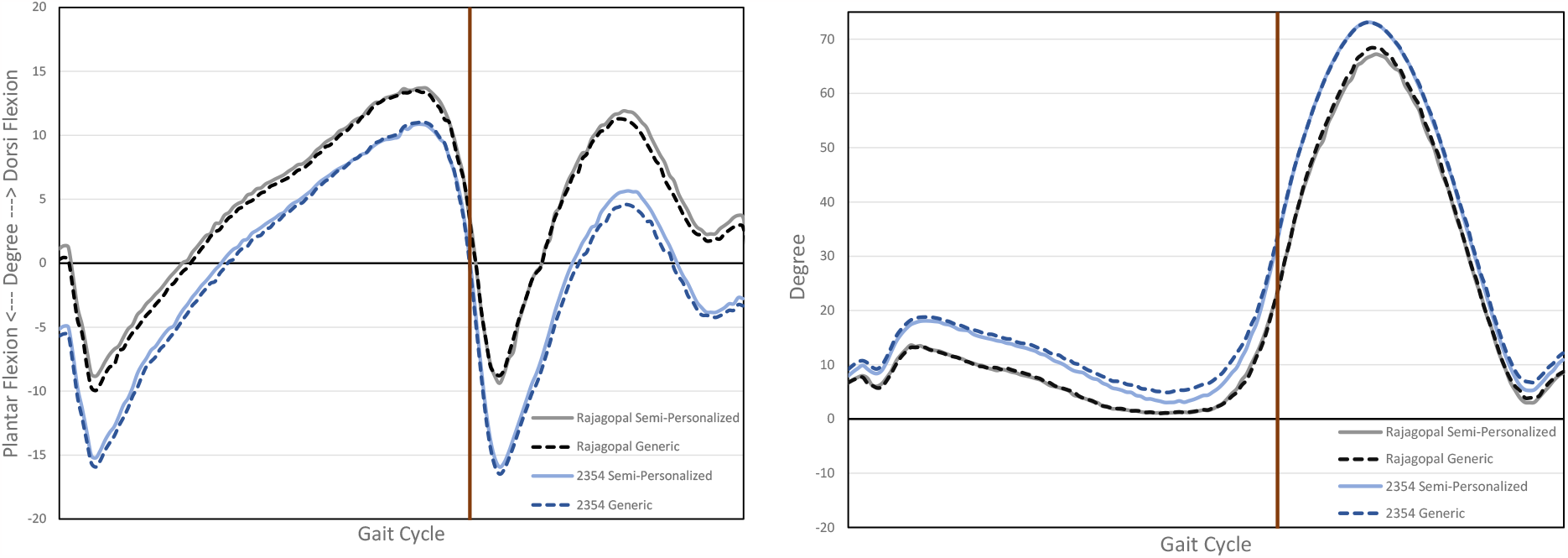
Ankle movement (right) in the sagittal plane and knee flexion(left) in semipersonalized and generic models of Gait 2354 and Rajagopal.

**Figure 6).**
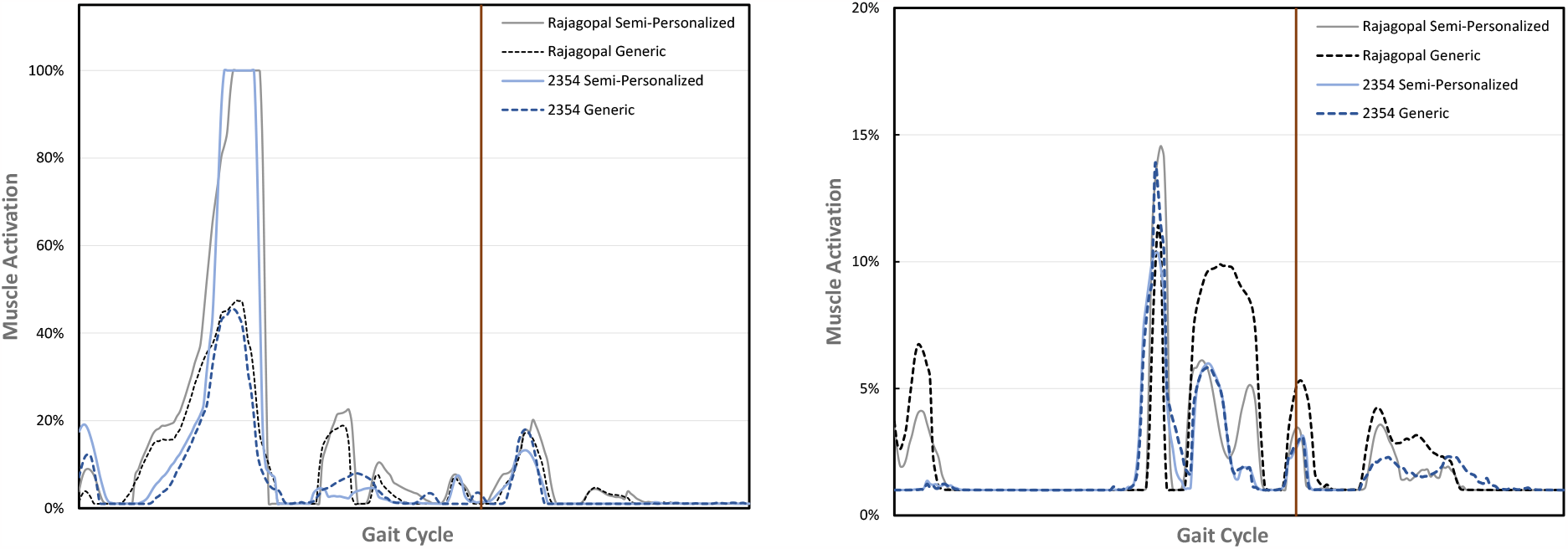
The activation pattern of Tensor Fasciae Latae (left graph) and Gracilis (right graph) for Rajagopal model and Gait 2354 per gait cycle.

To study the effects of bone deformation on muscles’ function, the muscle activation pattern for Tensor Fasciae Latae (TFL), a hip abductor, and Gracilis, a hip adductor, were estimated (shown in Figure 8). The activation pattern of TFL indicated a sharp rise during the stance phase of the semi-personalized model. On the other side, although the activation peak was seen for Gracilis in the semi-personalized model, the activation pattern has generally diminished.

## Discussion

The focus of this study was to develop and evaluate an OpenSim tool by which the user can modify the varus-valgus angulation of Rajagopal, Gait2354, and Gait 2392 models. The inputs of this tool are (1) an OpenSim model, (2) the magnitude of mMPTA, and (3) a CORA point, which is a virtual marker loaded in the model through OpenSim Graphical User Interface (GUI). Based on the magnitude of mMPTA and the location of CORA in the tibia, a deformation pattern is created. Muscles’ pathways, the tibia, and the location of virtual markers are manipulated using the deformation pattern, and eventually, a semi-personalized model will be created (Figure 1).

The first step of analyzing musculoskeletal systems in OpenSim is to scale the model to match the subject’s weight, height, and limb length. Since the bone geometries of generic OpenSim models are normal, the scaling process for patients with higher degrees of skeletal deformity can be complicated, time-consuming, and most importantly, inaccurate. This study proved that virtual and experimental markers are matched more precisely in semi-personalized than generic models. As a result, the maximum errors and root mean square errors deplete, which will raise the accuracy in higher-level analysis, like Computed Muscle Control (CMC).

After the scaling process, the model undergoes IK. As shown in Figure 4, most changes were detected in hip adduction/abduction movement. This is because of the fact that both varus/valgus angulation and hip adduction/abduction occur in the frontal plate. Changes in hip flexion/extension, hip internal/external rotation, knee flexion, and ankle dorsi/plantar flexion are also noticeable due to the decrease in maximum and RMS error in the scaling process and IK. Furthermore, this result supports Steif’s study (32), which demonstrated that varus malalignment is not an isolated problem in the frontal plane.

The reduction in markers’ maximum error in IK (Figure 3) and the significant differences in the results of paired-samples T-tests (Table 1 and Table 2) are the two noticeable signs that ensure that the semi-personalized model can effectively reduce the inaccuracies previously yielded invalid results.

Since the varus angulation causes some degrees of abduction in lower extremities, the study of hip or knee abductors and adductors becomes more critical. Mundermann et al. (33) summarized that hip abductor and adductor muscles could have a vital role in reducing knee adduction moment. Thus, the muscle activation pattern for TFL and Gracilis muscles were estimated (Figure 8). The activation of TFL increased after in semi-personalized model, which agrees with Lloyd’s conclusion (34), attributing the increase in TFL’s activation to the stabilization mechanism of the knee joint in varus deformity. In addition, the activation of hip abductors in the varus knee has a significant impact on stabilizing the pelvis and trunk in the frontal plane (35, 36), and eccentric activation of the knee abductors plays a critical role in the control of femoral angles in the coronal plane (37). On the other side, the activation range of Gracilis was lower in semi-personalized model than generic one although the activation peak of Gracilis after the modification was more than the generic one in the stance phase of the gait cycle. The decrease in Gracilis activation in the swing phase may be due to preparing the foot for initial contact while having varus deformity, which produces adduction angulation in the limb. In addition, since varus malalignment is considered as one of the leading causes of medial osteoarthritis of the knee, the reduced activation of adductor muscles, like Gracilis, could also be a brain strategy to slightly decrease the adduction moment in the knee joint, and therefore, postpone osteoarthritis in patients with varus malalignment.

One of the limitations of this study is the number of patients. The introduced tool was evaluated by only one subject. Thus, more subjects are needed to identify the possible problems in the tool. The other limitation is that the tool does not include any function to adjust inertial properties. It utilized the generic model’s inertial properties, which are not the exact properties of semi-personalized models. However, it would not be problematic in slight degrees of varus/valgus since changes in inertial properties are not significant. Additionally, inertial properties will be used in higher-level analyses in OpenSim, like ID and Static Optimization, and it would not affect results in the scaling and IK sections.

## Conclusion

In conclusion, personalizing tools are used to develop models which are ideally close to the personalized ones. Two distinct advantages of personalizing tools over personalized models are that they are time and cost-effective. Thus, the patient does not need to expose to any CT scan radiation. The varus/valgus tool introduced in this study can produce semi-personalised models by implementing varus/valgus angulation. Using this tool helps obtain more accurate and more realistic results, leading to more precise interpretation.

## Acknowledgement

The author(s) received no financial support for the research, authorship, and/or publication of this article.

